# Physical mechanisms of nanoparticle-membrane interactions: A coarse-grained study

**DOI:** 10.1101/2025.10.31.685850

**Authors:** Massimiliano Paesani, Ioana M. Ilie

## Abstract

Nanoparticles (NPs) are promising drug carriers for targeted therapies, diagnostic imaging and advanced vaccines. However, their clinical translation is limited by complex biological barriers that reduce cellular uptake and efficacy. Specifically, the interaction with the cellular membrane controls nanoparticle adhesion, wrapping or full engulfment, which ultimately govern nanoparticle internalization efficiency. Flexible nanocarriers (e.g. liposomes, polymeric, micelles) are particularly attractive because their deformability could help them enhance the probability of successful cellular entry. To understand the physical mechanisms associated with cellular uptake, we investigate the interaction of semi-flexible nanocarriers with a symmetric lipid bilayer using coarse-grained simulations. We represent a flexible nanoparticle using the previously introduced MetaParticle (MP) model and the membrane using the Cooke-Deserno model. By systematically varying nanoparticle properties, *i*.*e*., adhesion strength and topology, we identify distinct interaction regimes ranging from surface adhesion and trapping to complete wrapping and endocytosis. These regimes correlate with nanoparticle shape, size and surface properties, providing quantitative design principles for optimizing cellular uptake. Overall, this framework offers predictive insight into how the interplay between nanoparticle properties and membrane interaction governs cellular internalization, informing the rational design of next-generation soft nanocarriers.

## I. INTRODUCTION

Nanoparticles (NPs) have changed modern drug delivery strategies by offering better control over therapeutic payloads, enhanced targeting specificity and improved pharmacokinetics^1–6^. Flexible nanocarriers such as liposomes, polymeric nanogels and virus-like particles can be engineered to optimize drug release profiles, reduce off-target toxicity and navigate biological barriers^7,8^. Smartly tuning the mechanical properties of nanocarriers, whether soft or rigid, often proves decisive for circulation time, intracellular trafficking and successful cellular uptake^9^. Common nanocarrier development and optimization strategies focus on tuning surface functionalization, shape and size to enhance therapeutic efficacy and overcome biological barriers in complex physiological environments^10^.

Experimental approaches have yielded important insights, for example advanced precision-designed carriers^11^, flexible nanogel platforms^12^ and stimuli-responsive polymeric nanoparticle systems^13^, yet challenges remain in quantifying nanocarrier–membrane interaction mechanisms^14^. Computer simulations complement experiments by accessing molecular-scale mechanisms where resolution is insufficient. In particular, molecular dynamics (MD) simulations provided insights into the effects of nanoparticle surface functionalization on membrane insertion^15–17^. However, atomistic simulations of large processes, such as membrane encapsulation and uptake, are limited insufficient spatiotemporal reach due to prohibitive computational costs^18^. These challenges require the use of more efficient simulation strategies that can still preserve the essential physics and bulk properties of the systems. Coarsegrained (CG) models overcome the limitations of all-atom simulations by grouping multiple atoms into single interaction sites (“beads”), thereby reducing degrees of freedom and enabling simulations of larger systems over longer time- and lengthscales^19^. By preserving the relevant interactions, CG models can provide insight into the interaction mechanisms of nanoparticles with complex biological environments^20^. For instance, coarse-grained simulations have been fundamental in characterizing multifaceted effects-like as PEGylation^21^, antibody adsorption^22^, or ligand decoration^23^, providing insights into how variations in grafting density, polymer chain length and surface charge^24^ contribute to the overall mechanisms balanced during NP binding and membrane penetration. Detailed simulation studies have shown that the interplay between these factors can lead to differences in NP wrapping efficiency, the formation of pores and eventual membrane destabilization, which are all critical components of both endocytic and non-endocytic uptake mechanisms^25,26^. Furthermore, computational studies have successfully replicated experimentally observed trends in NP-induced membrane bending and provided mechanistic insights into the role of lipid rearrangements during NP adsorption and penetration^26–28^. Additionally, the interfacial properties of nanoparticles, such as ligand orientation, surface charge distribution and hydrophobicity, determine how NPs interact with cellular membranes. In particular, comparisons between advanced CG models and atomistic benchmarks showed that by fine-tuning the NPs characteristics lipid clustering, membrane deformation and NP wrapping ca be observed^29,30^. Such refinements include parameter adjustments that reproduce the energetic barriers associated with ligand translocation and maintain the fidelity of membrane deformation processes.

*In vitro* and *in vivo* studies have shown that the intrinsic flexibility of a nanocarrier regulates cellular uptake by enabling non-endocytic pathways and facilitating access to otherwise inaccessible targets^31^. Contrary, computational studies showed that increased particle softness enhances biocompatibility but reduces cellular uptake efficiency^32^. CG models typically treat nanoparticles as rigid bodies. While this representation can describe the properties of, for instance hard metallic nanoparticles, it does not capture the intrinsic flexibility of soft systems, such as lipid nanoparticles or even coverage with protein corona within the biological environment. As such, they can only partially reproduce the mechanical properties of the nanoparticles and do not capture the ability of a nanocarrier to adapt its shape, mechanical and interaction properties in response to the environment^33^. To overcome this challenge, we recently introduced the Metaparticle model, a coarse-grained model able to reproduce the intrinsic flexibility of nanoparticles^34^. In the Metaparticle representation, a nanoparticle is represented as a network of coarsegrained beads interconnected by springs, both with tuneable properties (Fig. 1, left). Such tunability can aid in understanding and controlling cellular uptake mechanisms of polymeric nanogels, liposomes and other soft-core platforms that can deform under physiological conditions^35^.

**FIG. 1.**
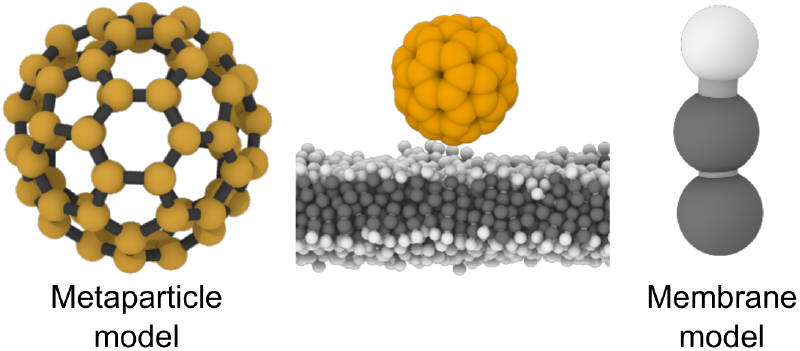
Coarse-grained model of the system. In a typical simulation, a flexible nanoparticle is placed on a lipid bilayer. Intrinsic particle flexibility is captured using the Metaparticle coarse-grained model^34^ (left), and membrane is modeled using the Cooke–Deserno coarse-grained model^36^ (right).

A bottleneck in the successful delivery of a cargo inside the cell, is the interaction of its nanocarrier with the cellular membrane. Here, we set the initial steps to understand the interaction mechanisms of flexible nanoparticles with the cellular membrane by using coarse-grained simulations. To this end, we use the metaparticle (MP) model^34^ to represent a deformable nanoparticle and study its interaction with a lipid bilayer, modeled via the Cooke-Deserno model^36,37^. By systematically varying MP-membrane affinity, we investigated how the metaparticle properties affect the membrane reshaping. We shed light on the interaction mechanisms between the MP and the membrane and studied the effects of nanoparticle size and topology on cellular uptake. Our results show that four distinct nanoparticle-membrane interaction regimes emerge: surface adhesion with negligible membrane stress and particle diffusion on the membrane surface, partial wrapping, full wrapping consistent with endocytic uptake and a kinetically trapped configuration that prevents successful internalization. Furthermore, our analysis shows that larger quasispherical nanoparticles are internalized more readily due to the favorable balance between adhesion energy and the membrane bending cost required for complete wrapping, whereas oblate particles remain trapped at partial wrapping states, consistent with Helfrich membrane elasticity theory. Finally, this work paves the way toward data-driven design rules for adaptive, topology-aware nanocarriers and bioinspired materials.

## II. SIMULATIONS DETAILS

Here, we focus on understanding the interaction mechanisms between flexible nanoparticles and the cellular membrane. We performed Langevin dynamics simulations, in the overdamped regime, of flexible nanoparticles with distinct sizes and topologies in proximity of coarse-grained lipid bilayers. For this, we combined a coarse-grained model for flexible nanoparticles (*i*.*e*., Metaparticle model^34^), which we recently introduced, with the coarse-grained phospholipid membrane model proposed by Cooke and Deserno^36,38^. In the Metaparticle model, a nanoparticle is represented as a collection of beads, interconnected by springs arranged in highly symmetric topologies (Fig. 1, left snapshot). In the Cooke model, phospholipids are represented by three beads, *i*.*e*., two hydrophobic and one hydrophilic, for the tails and the head, respectively (Fig. 1, right snapshot). To model the interaction between a metaparticle and the lipid membrane, we systematically varied the interaction between the metaparticle and the membrane beads. All the units used in this work are reduced Lennard-Jones units with *ε* and *σ* representing the units of energy and length, respectively. The time unit 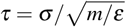 is expressed in terms of the particle mass, m, and the temperature given in *ε/k*_*B*_. A typical simulation system consists of a metaparticle and a homogenous coarse-grained membrane. The metaparticles used here are MP_60_, which consists of sixty beads displayed in an icosahedral arrangement^34^, the oblate MP_48_ and MP_180_. Assuming *σ* =1 nm, the effective diameters of the metaparticles are about 9 nm, 8 nm (along the longer axis), and 14 nm for MP_60_, MP_48_ and MP_180_, respectively, typical for small nanoparticles. The width of the membrane corresponds to about 5 nm. All simulations were carried out using the LAMMPS simulations package^39^, with an overdamped Langevin dynamics integrator in implicit solvent^40–43^ In preparation for the simulations, the metaparticle and the membrane were independently equilibrated to avoid artifacts. First, the metaparticle was equilibrated in a box (40×40×40 *σ*) with periodic boundary conditions for 10^5^ *τ* following the protocol we previously introduced^34^. Next, the membrane was first assembled in a hexagonal lattice to ensure periodicity. To reproduce a homogeneous membrane, 9472 lipids were arranged in a bilayer in a rectangular box (74×74×120 *σ*) with periodic boundary conditions. The membrane was equilibrated using overdamped Langevin dynamics in three steps alternating isoenthalpic–isobaric (NpH), micro-canonical (NVE) and NpH ensembles, each run for 5 · 10^6^ steps, to ensure membrane stability. This equilibration was run for 1.5 · 10^7^ steps in total, with a timestep equal to Δ*t* = 0.01*τ*. In the last step, the production run, the metaparticle was placed in the proximity of the relaxed membrane and simulations were run in the NpH ensemble for 4 million steps using a timestep Δ*t* = 0.01*τ* and at a fixed temperature of 1.1*ε/k*_*B*_. The potentials as well as the full simulation details can be found in the SI.

## III. RESULTS

### A. Metaparticle binding reshapes membrane topology

The simulations were started by placing the icosahedral MP_60_ in proximity of a symmetric lipid bilayer (Fig. 1). To understand the interaction mechanisms, we varied the attraction strength between the metaparticle and the membrane. Specifically, we systematically changed the affinity of the metaparticle beads (representing the functional groups on the surface of a nanocarrier) toward the lipid heads, *ε*_H_, and tails, *ε*_T_, between 2 and 9 *k*_*B*_*T* (corresponding to an effective affinity between −0.5 and −10 *k*_*B*_*T*, see Eq. S5 and Fig. S3).

For low affinity towards the lipid heads (*ε*_H_ ≤ 1), the metaparticle slides on the membrane surface and remains in contact over the duration of the simulation (Fig. 2(a)). Furthermore, the probability-density profiles of the lipids show a robust distribution of heads at ± 2.5 *σ* and tails around the center of the membrane. This indicates that the interaction with the nanoparticle, induces no structural changes in the membrane, highlighting that its structural integrity it is not altered (Fig. 2(a)). Increasing the affinity of MP_60_ towards the lipid heads (1 *< ε*_H_ *<* 4), drives membrane deformation and wrapping around the metaparticle (Fig. 2(b)). In this regime, the lipid heads gradually wrap around the metaparticle, reaching a maximum coverage of 45% (purple line in Fig. 2(b)). This partially wrapped state reflects a balance between adhesion energy from headgroup–metaparticle contacts and the membrane’s bending and tension penalties required for complete wrapping, consistent with continuum wrapping models^15^. The adhesion is sufficient to induce membrane deformation and invagination but insufficient to overcome the curvature and line-tension barriers needed for full wrapping and scission^15,44,45^. Furthermore, for tensionless bilayers, partially wrapped configurations can arise as thermodynamic equilibria, as shown in models that account for interactions in the neck region and flexible nanocarriers^32,46^. By increasing the affinity of MP_60_ towards the tails relative to the heads (*ε*_T_ *> ε*_H_), the number of tails around the metaparticle rapidly increases and stabilizes over the course of the simulations (yellow line in Fig. 2(c)). This indicates that the metaparticle inserts between the two leaflets of the bilayer and remains trapped over the duration of the simulation. Our results show, that upon insertion, the bilayer reorganizes, the lipids translocate and the hydrophobic tails turn toward the nanoparticle (left snapshot in Fig. 2(c)). Although this configuration is unlikely as an equilibrium state, it qualitatively captures how small hydrophobic nanoparticles can become kinetically trapped within membranes^47,48^.

**FIG. 2.**
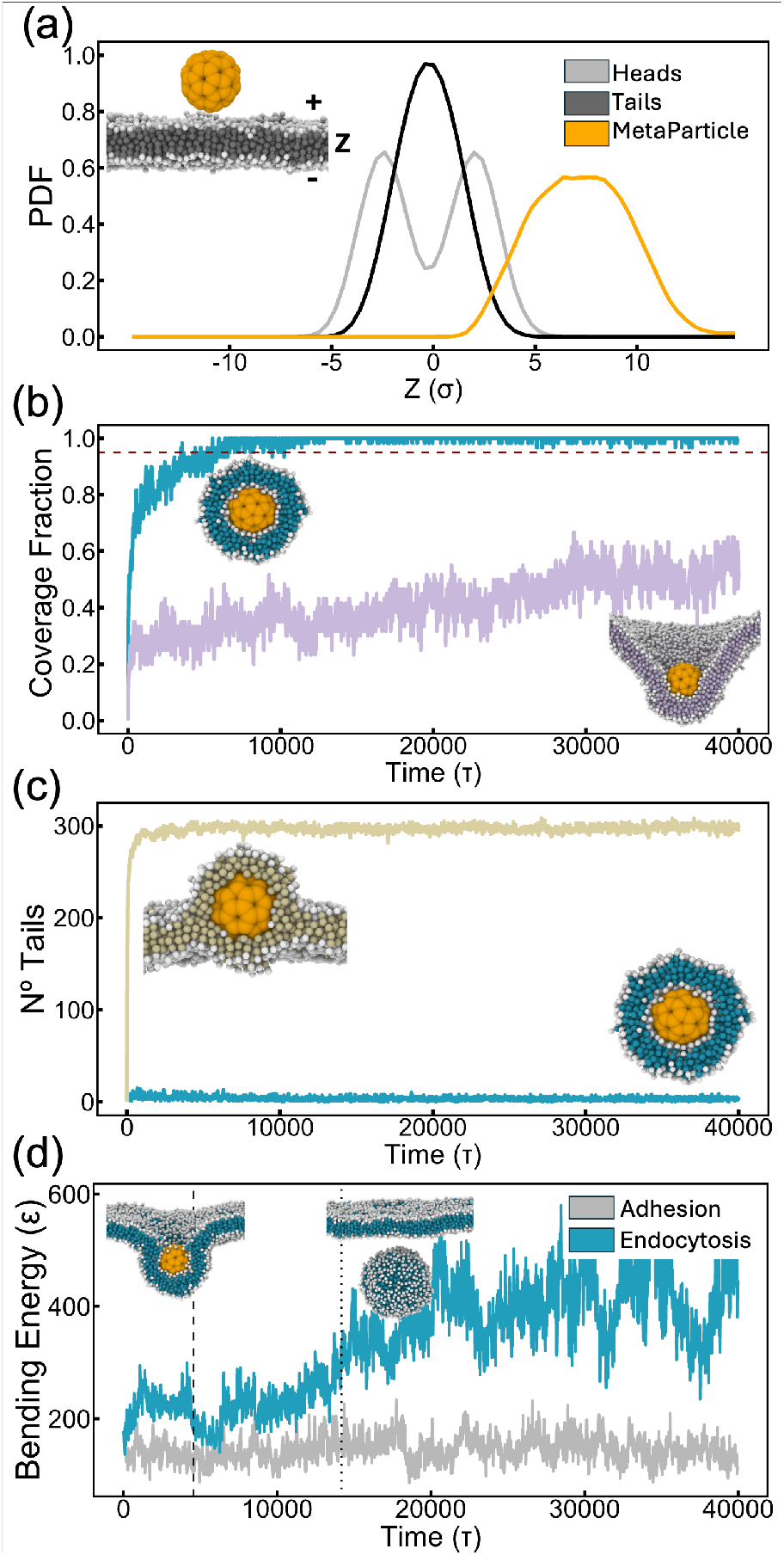
States identified in the simulations. (a) Probability distribution function of the lipid heads (grey), tails (black) and the MP_60_ beads (orange). (b) Timeline of the lipid head coverage fraction around the metaparticle. Compared are the profiles of a fully encapsulated (blue) and partially encapsulated MP_60_ (purple). The horizontal dashed line highlights 95 % coverage of the metaparticle. (c) Number of tail beads surrounding MP_60_ in the trapped state (beige) and the endocytosed state (blue) within a cutoff of 1.85*σ*. (d) Timelines of the membrane bending energies for for a metaparticle adhering to the membrane surface (grey) and an endocytosed MP_60_ (blue).

For *ε*_H_ *> ε*_T_, MP_60_ first adheres to the membrane, which then progressively wraps around the MP until it fully covers the MP with lipid heads (>95% coverage, (blue line in Fig. 2(b)), followed by the scission and the pinching off of the encapsulated bud (snapshots in Fig. 2(d)). The successful wrapping of the metaparticle results from the balance between adhesion and membrane bending^49^. Hence, we compared the bending energy of the membrane in the fully wrapped phase against the non-wrapped phase (adhesion). The blue profile in Fig. 2(d) shows the timeline of the bending energy calculated as explained in the SI. The profile shows an increase in bending energy as the the membrane wraps around the nanoparticle and subsequently pinch off from the rest of the membrane, reflecting the energetic cost of bending against membrane rigidity and tension. The profile shows a decrease in bending energy during the encapsulation pathway, which we ascribe to the reorganization of the membrane around the MP. As the contact area between the MP and the membrane increases during wrapping, *i*.*e*., as the coverage with heads increases (blue line Fig. 2(b)), the bending energy increases up to the point of full encapsulation and ultimately bud formation. Note that after the bud release, the bending energy remains high due to sustained curvature around the MP. In contrast, the non-wrapping membrane, shows lower and constant bending energy over the course of the simulation, indicating no curvature or strain induced by the MP (gray line Fig. 2(d)) As such, to achieve successful endocytosis, the adhesion energy must exceed membrane bending/tension costs^49^.

All in all, our results show, that depending on the conditions, four interaction regimes emerge for MP_60_ at the membrane: adhesion, partial wrapping, a kinetically trapped state and full endocytosis, each with a different membrane reshaping signature.

### B. Mechanisms and regimes

To understand the pathways associated with the four regimes, we performed k-means clustering. Briefly, k-means clustering is an unsupervised machine learning algorithm used for clustering unlabeled data by grouping similar items together around k representative “centers” (centroids)^50^. In practice, k-means relies on a distance metric in a numeric feature space to quantify the similarity between states. Here, the normalized feature vector comprises the position of the metaparticle center of mass along the membrane normal (z_MP_), the number of particles within a predefined membrane layer (N_mbr_), the local lipid-particle count surrounding the metaparticle (N_MP_) and the MP coverage fraction over time. From the identified centroids, we selected representative trajectories for further investigation. The analysis shows the temporal evolution of the MP (light to dark colors in Fig. 3(a)). MP tracking reveals that the metaparticles that occupy z_MP_ positions between –5*σ* and 5*σ* over the course of the simulation, reside predominantly near the membrane midplane, suggesting that they are kinetically trapped (beige line in Fig. 3(a)). Moreover, the MP coverage fraction discriminates between two different states, Fig2(b) and Fig.S5(b). One in which up to 50% wrapping of the membrane around the MP is reached, consistent with the partially wrapped state in Fig. 2(b), and one in which full MP coverage and detachment of the bud is reached, consistent with full endocytosis. Hence, a decreasing, and then stable, number of particles in the membrane layer suggest that the membrane wraps around the MetaParticle transiting towards an engulfed state (blue lines in Fig. 3(a), Fig.S5(a)). Fully encapsulated states are associated with a considerable increase (from 60 to 3000) and steady retention of particles surrounding the MP, and a pronounced reduction in the membrane slab (≈ 3000 particles)(Fig. S5(a)). Interestingly, the analysis revealed that the encapsulated MP primarily samples negative values along z_mbr_ and in isolated cases also positive values, suggesting bidirectional budding (Fig. 3(a) blue and dark blue lines, respectively). This outcome is consistent with a symmetric bilayer where fast encapsulation and local adhesion can trigger asymmetric lipid sorting, crowding or the emergence of curvature-preferring domains at the contact. This generates an effective spontaneous curvature that biases the direction of budding and scission without being influenced by the composition of the membrane leaflets^51^. Outward buddying can also be a kinetic pathway selection, particularly if the inward endocytic route faces higher barriers (*e*.*g*. neck formation), the system can relax along the outward budding pathway^52^. Interestingly, our analysis revealed a budded MP that follows the inward endocytic pathway, but contains a reduced number of particles (≈ 1150 as compared to the others) in the formed vesicle. Visual inspection, revealed that the nanoparticle was first adsorbed between the lipid leaflets being exclusively surrounded by lipid tails and subsequently the bud was released by the membrane. We attribute this to a spontaneous membrane bending, that facilitated the release despite tail-only contacts. Single-layer assemblies such as lipoproteins exist in nature^53,54^ and tail-facing monolayers can be engineered for certain nanoparticle insertions^55,56^. However, to our knowledge, there is no experimental evidence that a tail-only monolayer can form through a complete endocytic process.

**FIG. 3.**
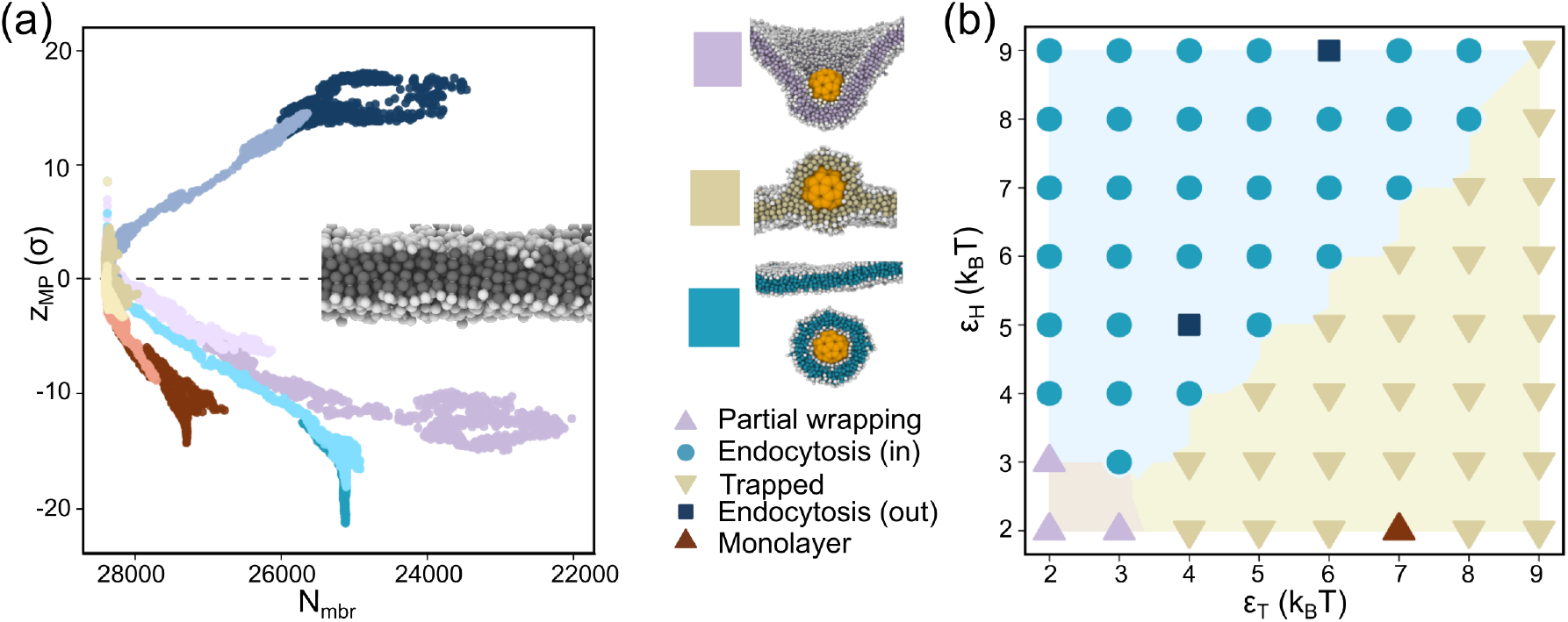
Uptake pathways and phase diagram. (a) Two-dimensional projection of representative simulation timelines for each identified regime. Shown are the center-of-mass position of MP_60_ along the membrane normal (z_MP_) with 0 the bilayer midplane versus the number of membrane-beads in a layer of 17 *σ* centered on the bilayer midplane (N_mbr_). The simulation time is shown by the color intensity, evolving from lighter to darker colors. Six different scenarios for MP-membrane interaction are identified: trapped (beige), partial wrapping (purple), enclosed by a bilayer membrane and endocytosed inward (blue) or outward (dark blue), and internalized with a monolayer wrapped around the MP (brown). (b) Phase diagram showing the uptake outcomes depending on the affinity of the metaparticle beads toward the lipid head (*ε*_H_) and tails (*ε*_T_).

To better understand the emerging regimes, we projected the k-means clustering onto a phase diagram with *ε*_H_ and *ε*_T_ along the axes. (Fig. 3(b)). The analysis reveals that states segregate along the diagonal component of the diagram, with the upper triangular region corresponding to MP encapsulation within a bilayer and the lower triangular region capturing trapped states. This partition highlights a transition boundary between wrapping and arrest, which could be exploited in the future design of nanoparticles to either promote membrane wrapping or achieve stable trapping within the bilayer.

### C. Endocytosis set by size, topology and adhesion thresholds

To investigate how size and geometry influence uptake, we performed an additional set of simulations for MP_180_ and MP_48_. MP_180_ has icosahedral symmetry but is larger in diameter (≈ 14 nm) than MP_60_, enabling studying size effects on cellular uptake. MP_48_ is slightly smaller than MP_60_ in diameter (≈ 8 nm) but exhibits D_6d_ symmetry and, according to our previous study, presents the highest toughness among the tested metaparticles^34^.

Upon interaction with the membrane, MP_60_ adheres to the bilayer (t = 0 *τ*) (Fig. 4(a)). Subsequently, the membrane wraps around the MP, reaching 50% coverage within 200 *τ*. Full nanoparticle wrapping requires longer times and is achieved within the next ≈ 10.000 *τ*. The pinching off of the bud requires almost half as much time due to the additional barriers associated with neck scission. The larger MP_180_ adheres to the lipid bilayer in a similar fashion (t = 0 *τ*) prior to inducing membrane deformation (Fig. 4(b)). The wrapping smoothly continues as the contact area grows and 50% of the nanoparticle is covered by lipid heads within the next 740 *τ*. After 17000 *τ*, the metaparticle is fully covered by the lipid bilayer and the bud pinches off within the subsequent 1000 *τ*. Qualitatively, both metaparticles follow the same encapsulation pathway, however the larger MP requires a larger wrapping area and increased membrane bending before neck closure. Variations in adhesion strength modulate the underlying energy barriers, hence the observed timescales are not directly comparable across conditions. Upon initial adhesion (t = 0 *τ*), MP_48_ rotates on the membrane surface to expose its oblate side, thereby increasing contact area and adhesion (Fig. 4(c)). The membrane then wraps around the MP to reach 50 % coverage within 560 *τ*, increasing the surrounding membrane tension. Subsequently, the MP rotates vertically while maximizing its adhesion (around ≈ 20000*τ*), though no closure of the neck or budding occurs on the timescale of the simulation. Because adhesion and curvature stresses do not overcome a critical threshold, MP_48_ remains trapped in a deeply wrapped, non-budded state^57^.

**FIG. 4.**
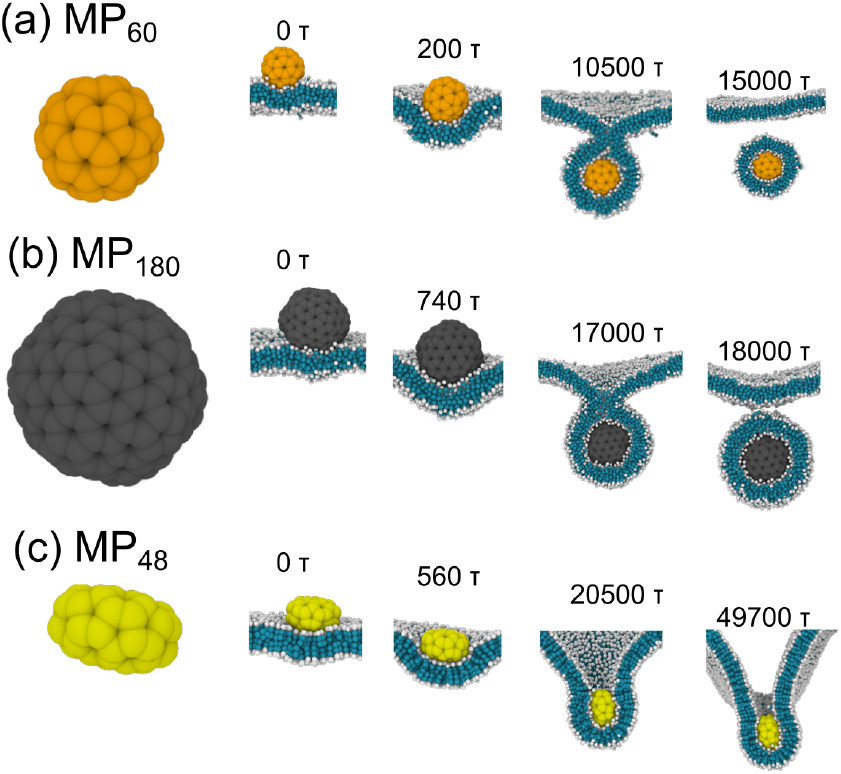
Endocytosis of nanoparticles of different sizes and topologies. Compared are (a) MP_60_ for *ε*_H_ = *ε*_T_ = 3*k*_*B*_*T*, (b) MP_180_ for *ε*_H_ = *ε*_T_ = 2*k*_*B*_*T* and (c) for *ε*_H_ = *ε*_T_ = 3*k*_*B*_*T*. The metaparticles adhere to the membrane and reach 50% membrane coverage within the initial 1000*τ*. MP_60_ and MP_180_ are fully wrapped and undergo endocytosis, while MP_48_ is kinetically trapped.

To compare against existing theories, we assume MP_60_ and MP_180_ behave like spheres with effective membrane interaction radii of ≈ 4.5 nm and ≈ 7 nm, respectively. From the Helfrich theory of membrane elasticity^58^, the bending energy for a sphere corresponds to 8*πκ*, with *κ* the bending rigidity of the membrane associated with mean curvature independently of size, assuming no spontaneous membrane curvature and zero membrane tension. The condition of full wrapping is for the adhesion energy gained by increasing contact area to exceed the elastic costs of bending and membrane tension^49^, which results for the radius to be larger than 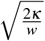, with w the adhesion energy per unit area. Hence, smaller particles require stronger adhesion per unit area to overcome membrane bending resistance. Concretely, for MP_60_ an effective attraction strength of 0.5 *k*_*B*_*T* between the MP beads and lipid heads (corresponding to *ε*_H_ = 2 *k*_*B*_*T*, see Fig. S3 and Eq. S5) results in an adhesion energy per unit area of 0.2 *k*_*B*_*T/*nm^2^ (MP_60_ surface area of 139 nm^2^ and *κ* = 10 *k*_*B*_*T*), which is sufficient to enable successful endocytosis for a spherical nanoparticle with a diameter of 9 nm. Increasing the particle radius, and implicitly the available surface area (≈ 335 nm^2^), while preserving the MP topology as in the case of MP_180_, enables full encapsulation at lower effective interaction strength (*ε*_H_ = 2*k*_*B*_*T*), corresponding to an increased adhesion energy w = 0.3 *k*_*B*_*T/*nm^2^. This translates to a minimum wrapping diameter of about 8.6 nm, consistent with the observed successful wrapping of the larger nanoparticle, which was not achieved for MP_60_. Hence, our results show that increasing the radius of the metaparticle while preserving the topology, wrapping becomes thermodynamically more favorable as the critical adhesion energy per unit scales with the radius of the particle. MP_48_ has D_6d_ topology and resembles a flattened spheroid, having low curvature and high curvature regions. In our simulations, MP_48_ binds to the membrane at its low-curvature face to maximize adhesion and initiate wrapping. As the lipid heads reach the high-curvature face, the membrane bending cost increases, which drives the reorientation of the metaparticle to preserve favorable curvature matching. The neck does not close because adhesion is insufficient to overcome the curvature-dependent scission barrier, leaving MP_48_ in a kinetically trapped state. Endocytosis would become energetically possible when the adhesion energy overcomes the bending penalty (Eq. S12).

## IV. DISCUSSION AND CONCLUSION

The efficacy of a nanocarrier is linked to its ability to overcome biological barriers and deliver its cargo inside the cell. A bottleneck in the successful transport of nanoparticles to the desired target is the interaction with the cell membrane. Essentially, the membrane acts as a protective layer for the cell, which controls the passage of macromolecules and nanoparticles, and its disruption can lead to fatal consequences^59,60^. Here, we investigated the physical mechanisms of interaction between flexible nanoparticles and a lipid bilayer. For this, we use the metapaticle coarse-grained model for flexible nanoparticles of different sizes and topologies^34^ combined with the Cooke-Deserno model for fluid membranes^36^.

To understand nanoparticle-membrane interaction mechanisms, *i*.*e*., to probe adhesion, specificity and receptor effects, we tuned the effective interactions between metaparticle beads and the lipid head- and tail-group beads. Our results show that MP binding reshapes membrane topology, which leads to four critical regimes, *i*.*e*., adhesion, partial wrapping, trapping and endocytic encapsulation, consistent with experimental results and observations. First, at low MP-membrane affinity, the metaparticle adsorbs and diffuses on the bilayer surface without inducing any membrane deformation. In biological systems, a protein corona or adsorbed surfactants can modify the surface properties of the nanocarrier thereby changing their effective interaction with the membrane. As such, protein coronas were shown to reduce membrane binding and implictily cellular uptake^61,62^, while surfactant coatings weakened the adhesion to the membrane^63^. Second, at moderate affinity towards the lipid heads, the metaparticle is in a partially wrapped state, remaining about 50 % covered by lipid heads. This trapped state often arises when the adhesion energy is sufficient to induce membrane deformation but insufficient to overcome the bending cost of encapsulation^64,65^ and has been previously observed in the case of flexible nanocarriers^32,66^. Third, when the attraction to lipid heads is sufficiently strong, the MP drives the membrane deformation, becomes fully wrapped and subsequently pinches off from the membrane to undergo a full endocytic process. This is in line with both experiments and simulations across different scales, conditions and nanocarrier surface properties^67–71^. Fourth, when the attraction to the lipid is strong, the nanoparticle rapidly becomes fully embedded in the membrane and remains trapped inside the lipid bilayer throughout the simulations. This kinetically trapped state replicates the behavior of small hydrophobic nanocarries that can embed in the bilayer^47^ or can induce lipid translocation^48^. These trapped stated can be controlled in context of sensing and imaging or even be used to induce membrane disruption for immunotherapy^72–75^. Taken together, these results underline the importance of engineering nanocarriers not only for initial adhesion and uptake, but also for corona management, with quantitative thresholds for wrapping and escape guiding the design space.

Next, we investigated how the nanoparticle properties (e.g. size, geometry, membrane affinity) affect endocytosis, from adhesion to the membrane and wrapping to neck closure and detachment, using quasi-spherical and oblate metaparticle morphologies (MP_180_ and MP_48_). Our results show that preserving the particle topology while increasing its diameter enables successful endocytosis, which can occur even at a lower adhesion energy per unit surface area. This is because a larger MP provides a higher contact area with the membrane, thereby increasing the overall adhesion energy required to overcome the bending resistance required for full wrapping^26,59,64^. In contrast, the oblate MP_48_ fails to achieve full encapsulation and remains trapped in a deeply partially wrapped state, in which the adhesion energy is insuffucient to overcome the membrane curvature, consistent with theoretical predictions^26,59,64^. This observation agrees with previous theoretical and computational studies showing that highly oblate or disk-like nanoparticles experience unfavorable curvature matching and often arrest in metastable, partially wrapped states due to insufficient adhesive gain relative to bending energy costs^20,46,49,76–78^. Internalizing an oblate nanoparticle requires the cell membrane to bend and curve around a flat surface with sharp edges, which is energetically expensive. Biologically, this means that the cell must spend more energy (and possibly recruit more receptors, cytoskeletal forces) to wrap a system like the oblate MP_48_ completely^78–80^, as compared to wrapping a spherical system of similar surface area like MP_60_. Importantly, the initial nanoparticle orientation relative to the membrane controls how the process unfolds^28^. To be internalized, an oblate structure has to interact through the shortest cross-section initially, rather than lying flat on the longer side^79–81^. Approaching the membrane in this way, the nanoparticle would expose the smaller contact area, which can lower the energetic barrier for the membrane to start curving around it^79–81^.

Our simulation results provide guiding principles for the rational design of flexible nanocarriers. For instance, coupling metaparticle dynamics with free-energy calculations can provide quantitative insight into the kinetic barriers associated membrane wrapping^82,83^ or thermodynamics of pore formation^84–86^. Integrating the MP framework with machinelearning–driven optimization and inverse-design strategies further enables data-guided discovery of nanoparticle architectures and properties that control cellular uptake and bilayer trapping^87,88^. From a design perspective, an emerging paradigm is mechanotargeting, which emphasizes optimizing nanoparticles for their mechanical interactions with cellular membranes^89^. Flexible nanocarriers represented through the metaparticle model can be engineered to respond in a controlled fashion to external stress or variations in their local microenvironment. This adaptive mechanical behavior can be exploited in context of efficient membrane wrapping and direct cytosolic release. In the current model, we envision a dynamic regime in which the metaparticle beads and the spring network can adjust their properties in response to the environment. For instance, the spring stiffness can be adjusted depending on the proximity to or embedding in the membrane, hence capturing MP mechanical response to membrane-induced stress. Furthermore, bead properties can be altered to investigate the effects of multivalency on cellular uptake^90^. Such efforts can then be used to derive fundamental design principles for next-generation nanocarriers that replicate the adaptive mechanics of natural systems, including viruses and bacterial vesicles^31,91,92^. This is relevant for drug delivery, since nanocarriers with tuned properties may lower free energy barriers to release their payload directly into the cytosol, bypassing endosomal degradation^9,93^. Moreover, recent peptide-based nanoparticles show the impact of topology on nanocarrier properties and functions, futher underscoring the applicability of metaparticles as a model system for exploring topology-driven design principles^94^. Overall, the significance of this work can be extended beyond drug delivery, where it provides a foundation for linking particle shape, flexibility, multivalency and membrane mechanics to predictable uptake and release, and into the rational design of adaptable, bioinspired materials and the study of nanomaterial interactions with elastic or solid interfaces.

## ACKNOWLEDGMENTS

I.M.I. acknowledges support from the Sectorplan Bèta & Techniek of the Dutch Government, the Dementia Research - Synapsis Foundation Switzerland and the Molecular Material Design Technology Hub.

## V. DATA AVAILABILITY

The input files are publicly available on: https://github.com/ilieim/NanoparticleMembrane

