## Supplementary Information for "Physical mechanisms of nanoparticle-membrane interactions: A coarse-grained study"

### I. MODEL AND METHODS

#### A. MetaParticle model

To represent a flexible nanoparticle, we use the previously introduced *MetaParticle* model[1]. A metaparticle consists of a collection of beads, interconnected by springs arranged in highly symmetric topologies (Fig. S1)

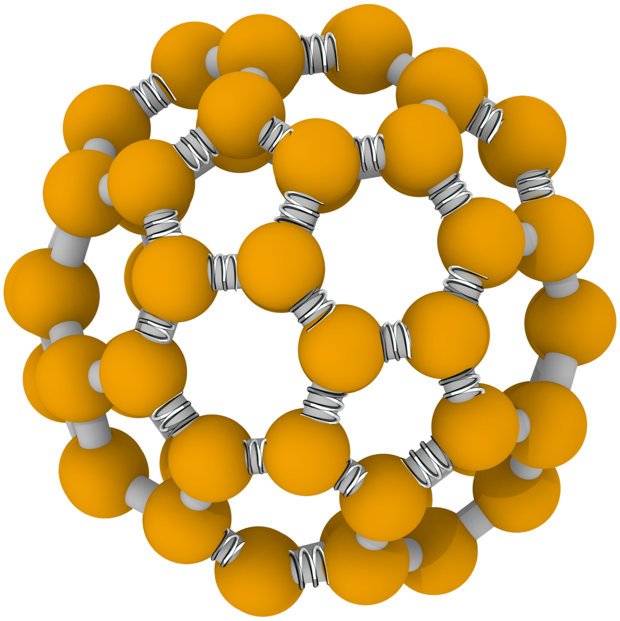

Fig. S1. A MetaParticle $MP_{60}$ that consists ofsixty60 beads interconnected by ninty bonds in a symmetric arrangement.

The interactions between the beads are described by bonded and non-bonded interactions between interconnected and non-connected beads, respectively. The non-bonded interactions are captured by the Lennard-Jones potential:

$$\Phi_{\mathrm{NB}}^{\mathrm{MP}}(r) = \begin{cases} 4\,\varepsilon_{\mathrm{LJ}}\left[\left(\frac{\bar{\sigma}}{r}\right)^{12} - \left(\frac{\bar{\sigma}}{r}\right)^{6}\right], & r < r_c, \\ 0, & r \geq r_c\,. \end{cases} \tag{S1}$$

where $r$ is the distance between the centers of mass of the beads, $\bar{\sigma}$ the particle diameter and $\varepsilon_{\mathrm{LJ}}$ the interaction strength. The bonded term, accounts for the connectivity between the particles while preventing overlap, and is described by

$$\Phi^{\mathrm{MP}}_{\mathrm{bond}}(r) = -\frac{1}{2}Kr_0^2 \log\left[1-\left(\frac{r}{r_0}\right)^2\right] + \Phi^{\mathrm{MP}}_{\mathrm{WCA}} \qquad \text{for} \quad 0 < r < r_0 \tag{S2}$$

where $r_0 = 1.8\sigma$ is the maximum bond extension, K=30 $\varepsilon/\sigma^2$ the spring constant of the potential and the repulsive potential $\Phi^{\mathrm{MP}}_{\mathrm{WCA}} = 4\varepsilon\left[\left(\frac{\sigma}{r}\right)^{12} - \left(\frac{\sigma}{r}\right)^{6} + \frac{1}{4}\right]$ for $r < 1.12\sigma$.

#### B. Membrane model

To define the biological membrane, we employed a coarse-grained model as introduced by Cooke and Deserno[2]. This membrane model is known to be able to capture wrapping, fusion and direct translocation events[3,4]. In the Cooke model each lipid is represented as a set of three beads, one hydrophilic and two hydrophobic, for the head and the tails, respectively (Fig. 1). The interaction between the beads is described by bonded and non-bonded potentials. The bonded interactions within a lipid consist of a spring-like FENE potential and a WCA potential to account for the connectivity between the beads and prevent unphysical streching, and a bending potential to control the stiffness of the lipid tails $\Phi^{\mathrm{mbr}}_{\mathrm{bend}}$. The non-bonded interactions control the membrane assembly and fluidity and are represented by a hydrophobic potential, $\Phi^{\mathrm{mbr}}_{\mathrm{cos}}$.

As the first two have the same expression as Eq. (S1) we will focus on introducing only the bending and the non-bonded potentials. The bending term controls the flexibility of the lipid tail via the harmoninc angular potential

$$\Phi^{\mathrm{mbr}}_{\mathrm{bend}}(r) = \frac{1}{2}k_{\mathrm{a}}\left(r - 4\bar{\sigma}\right)^2, \tag{S3}$$

where $k_a = 10\varepsilon$ is the bending stiffness, $r$ the distance between two beads and $\bar{\sigma}$ the effective interaction diameter.

The hydrophobic term is critical for recovering the fluidity of the membrane bilayer and is described by an attractive cosine-based potential[2,5,6]

$$\Phi^{\mathrm{mbr}}_{\mathrm{cos}}(r) = \begin{cases} -\varepsilon_{\cos} & r < r_c \\ -\varepsilon_{\cos}\cos^2\left[\dfrac{\pi(r-r_c)}{2w_c}\right] & r_c \le r \le r_c + w_c \\ 0 & r > r_c + w_c, \end{cases} \tag{S4}$$

where $\varepsilon_{\cos}$ the depth of the potential, which smoothly decays to zero beyond the cutoff distance $r_c$. The stiffness and fluidity of the membrane is adjusted by tuning the parameter $w_c$ to reflect membrane fluidity. For the FENE potential, the spring constant was set to 30 $(\varepsilon/\sigma^2)$ and the maximum bond length $r_0$ equal to $1.5\sigma$. In the simulations the effective interaction diameter was set to $\bar{\sigma}_{\text{head-head}} = \text{head-tail} = 0.95\sigma$ and $\bar{\sigma}_{\text{tail-tail}} = \sigma$ to ensure the cylindrical shape of the lipid molecules and the We set $w_c = 1.6\sigma$, the value identified by Cooke–Deserno as optimal for a stable fluid-phase membrane[2].

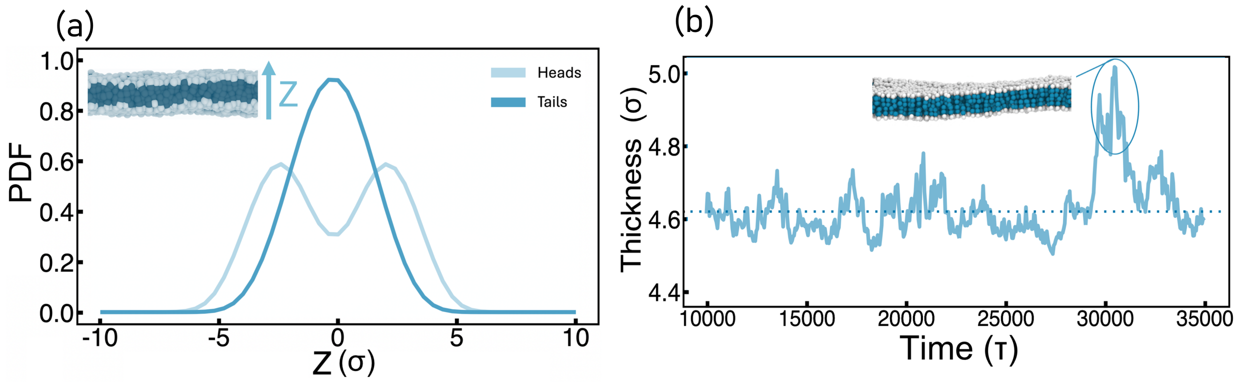


Fig. S2. (a) Probability distribution of lipid head and tail beads along the membrane normal ($z$-axis). The head- and tail-beads profiles confirm the stablity of the bilayer over the course of the simulations. (b) Membrane thickness as a function of time, showing minimal fluctuations and indicating that the membrane remains structurally stable over the course of the simulation.

#### C. Metaparticle-membrane Interaction Potentials

The interaction between a metaparticle and the membrane is described by a 12-6 Lennard-Jones potential, in which the Lorentz-Berthelot rules[7] to determine the effective interaction diameter between the beads in the metaparticle and the membrane beads.

$$\Phi_{\text{NB}}^{\text{MP-mbr}}(r) = \begin{cases} \mu(r,\bar{\sigma}) \cdot \left[\varepsilon_{\text{rep}} \left(\frac{\bar{\sigma}}{r}\right)^{12} - \varepsilon_{\text{attr}} \left(\frac{\bar{\sigma}}{r}\right)^{6}\right] & \text{if } r < 2.5\bar{\sigma} \\ 0 & \text{if } r \geq 2.5\bar{\sigma} \end{cases} \tag{S5}$$

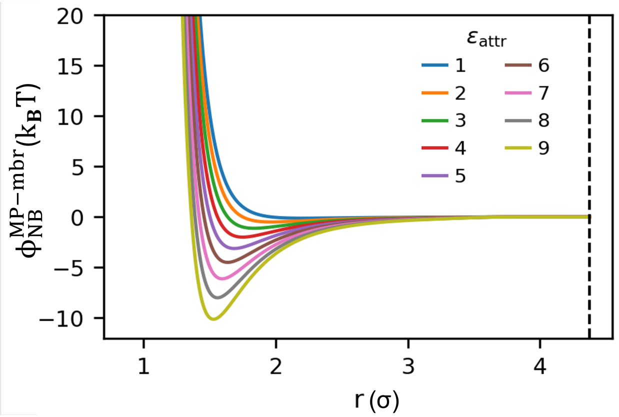


Fig. S3. Interaction potential between the metaparticle and the membrane components (heads and tails of the lipids) for different attractive strengths. The effective interaction length $\bar{\sigma} = 1.75\sigma$, which represents the interaction between the metaparticle and the lipidic tails, with $\alpha = 5$ and the cut-off distance set to $2.5\sigma$.

where $\mu(r)$ is a step function that smoothly decays from 1 to 0 to ensure the continuity in the potential and the forces, while keeping the repulsive part constant independently of $\varepsilon_{\text{attr}}$.

$$\mu(r, \bar{\sigma}) = \frac{1}{2}\left[1 - \frac{\tanh\big(\alpha(r - 2\bar{\sigma})\big)}{\tanh\big(2\alpha\bar{\sigma}\big)}\right]. \tag{S6}$$

where $\alpha$ determines the width of the transition region. We set the repulsive interaction strength to $\varepsilon_{\text{rep}} = 2k_BT$ to model a repulsive soft metaparticle. The attractive interaction strength $\varepsilon_{\text{attr}}$ was varied from 2 to 9 across different simulations (64 in total) to probe the effect of varying binding affinities with membrane receptors. Specifically, $\varepsilon_{\text{attr}} = \varepsilon_{\text{H}}$ denotes the affinity toward the lipid heads, while $\varepsilon_{\text{attr}} = \varepsilon_{\text{T}}$ corresponds to the interaction with the lipid tails. The sharpness of the potential transition was controlled by the parameter $\alpha = 5$. The effective interaction diameter $\bar{\sigma}$ was determined according to the Lorentz–Berthelot mixing rules[7], ensuring consistency between the metaparticle beads and the membrane head or tail beads.

### II. K-MEANS CLUSTERING

To discriminate between the emerging states, we employed k-means clustering. We feed the algorithm with four (normalized) quantities tracked over time: $\text{N}_{\text{mbr}}$, which represents the total number of particles inside the membrane, $\text{N}_{\text{MP}}$, which represents the number of membrane particles around the metaparticle, $z_{\text{MP}}$ which represents the MP center of mass position along the z

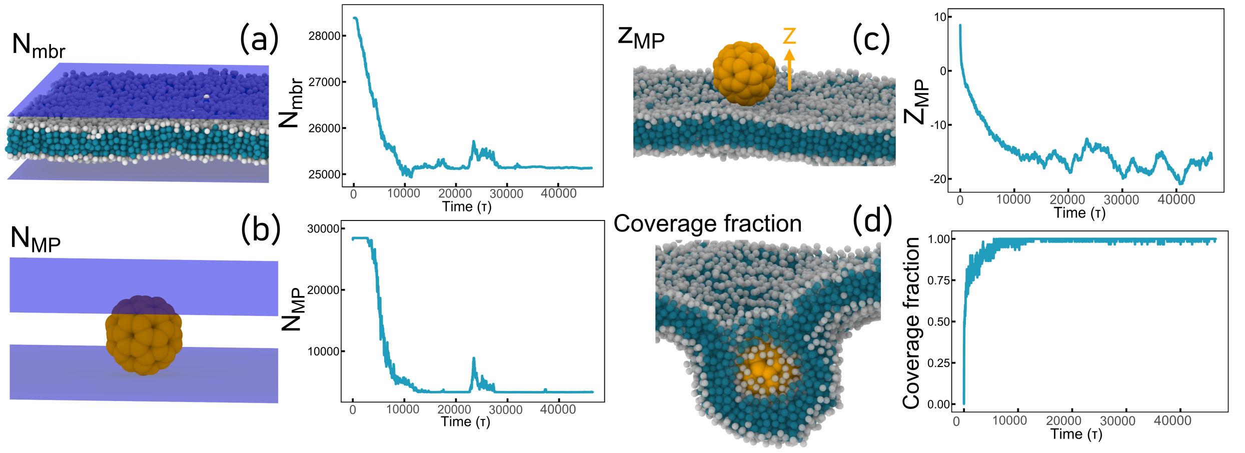


Fig. S4. Data input for the *k*-means clustering analysis. (a) We defined a 17$\sigma$-thick layer at the membrane center of mass and record the number of particle inside it over time. (b) We similarly define a $\approx 20\sigma$-thick layer centered on the metaparticle center of mass and track the number of particles within it. (c) We record the metaparticle center-of-mass position along the $z$-axis over time. (d) We compute the metaparticle coverage fraction, defined as the fraction of its surface area in contact with lipid beads, and track its evolution throughout the simulation.

axis over time, and lastly the coverage fraction, which represents how much the metaparticle is surrounded by lipidic particles within 1.85$\sigma$ from the MP center of mass. We selected four centroids, each corresponding to a representative state within the phase diagram, enabling the algorithm to discriminate configurations according to these reference points. From the clustering results, we identify six distinct states: (i) partial wrapping, where the metaparticle remains wrapped but cannot complete membrane scission; (ii) endocytosis, where the metaparticle becomes fully encapsulated and detaches from the membrane; (iii) trapped, in which the metaparticle is kinetically arrested within the bilayer; (iv) a monolayer-associated state, observed for highly hydrophobic metaparticles, where wrapping occurs only on one leaflet; and (v–vi) upper-leaflet endocytosis, reflecting symmetry of the bilayer and occurring when internalization proceeds from the opposite membrane side. The states were all discriminate correctly but the one we refer as ’monolayer’, was discovered upon visual inspection of all the trajectories.

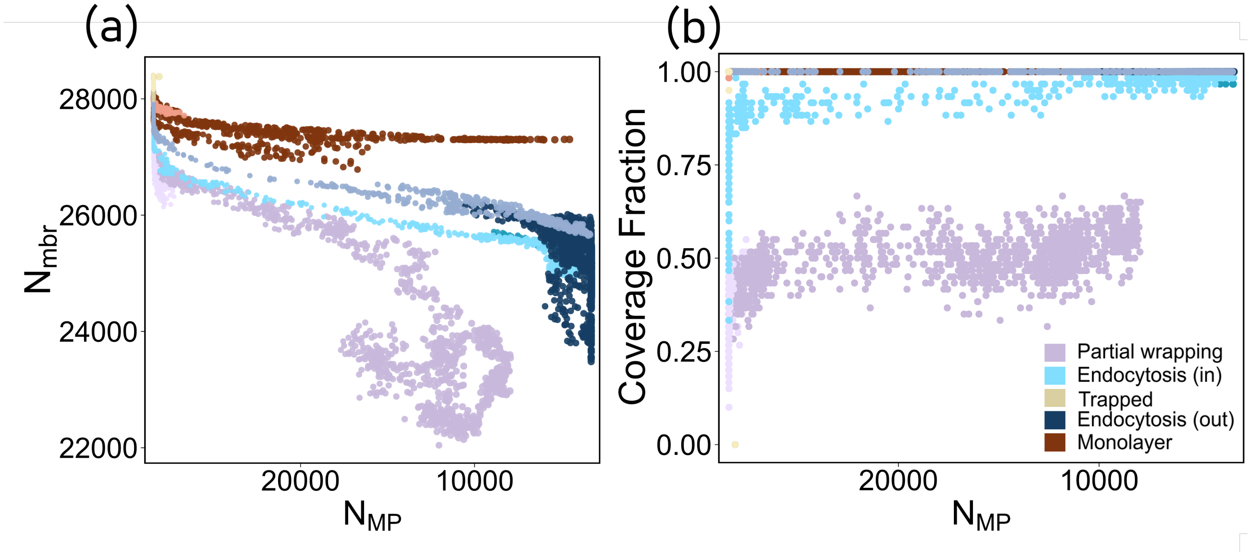


Fig. S5. (a) Number of metaparticle beads within the membrane-centered layer plotted against the number of beads within the metaparticle-centered layer. Partially wrapped states consistently show a high number of beads in the metaparticle layer, whereas the number of beads in the membrane layer decreases due to the strong local membrane curvature, which biases the layer-based counting. (b) MP coverage fraction. Partially wrapped states plateau at approximately 50%, indicating incomplete encapsulation.

### III. BENDING ENERGY

For a tensionless membrane with no spontaneous curvature, the free energy of local membrane deformation can be expressed via the Helfrich equation

$$F = \int \frac{\kappa}{2}(c_1 + c_2)dA + \int \frac{\bar{\kappa}}{2}(c_1 c_2)^2 dA \qquad \text{(S7)}$$

where $c_1$, $c_2$ the local principal curvatures of the membrane, $\kappa$ and $\bar{\kappa}$ the bending moduli of the membrane associated with the mean and Gaussian curvatures, respectively. For the subsequent steps, we will assume the membrane to have a closed topology and set $\bar{\kappa} = 0$.

#### A. Sphere

The surface area of a sphere with radius $R$ is $A = 4\pi R^2$, which leads to a bending energy $E_B = 8\pi\kappa$ independently of the particle radius. Full encapsulation becomes thermodynamically favorable when the adhesion to the membrane balances out the bending costs of the membrane, *i.e.,* when the total change in free energy is negative, $\Delta F = E_B + E_A \leq 0$, with $E_A = -wA$ the adhesion energy. For a sphere, this gives a minimum radius

$$R_{\mathrm{min}} = \sqrt{\frac{2\kappa}{w_c}}. \tag{S8}$$

#### B. Oblate ellipsoid

For an oblate ellipsoid with the principal axis components $c < a$, the surface area is[8]

$$A = 2\pi a^2 + \frac{\pi a c^2}{\sqrt{a^2 - c^2}}\lambda, \tag{S9}$$

with $\lambda = \ln\left(\frac{a+\sqrt{a^2-c^2}}{a-\sqrt{a^2-c^2}}\right)$. Assuming a tensionless membrane with no spontaneous curvature, the Helfrich bending energy expressed only in terms of the mean-curvature becomes

$$E_{\mathrm{bend}} = \pi\kappa\left[\frac{14}{3} + \frac{4a^2}{3c^2} + \frac{c^2}{a\sqrt{a^2-c^2}}\lambda\right]. \tag{S10}$$

One can readily determine the adhesion energy for full wrapping as

$$E_{\mathrm{adh}} = -wA = -w\left(2\pi a^2 + \frac{\pi a c^2}{\sqrt{a^2-c^2}}\lambda\right), \tag{S11}$$

which results in

$$\frac{w_c}{\kappa} = \frac{\frac{14}{3} + \frac{4a^2}{3c^2} + \frac{c^2}{a\sqrt{a^2-c^2}}\lambda}{2a^2 + \frac{ac^2}{\sqrt{a^2-c^2}}\lambda}. \tag{S12}$$

For a sphere of $c = a$ the expression reduces to $w_c = 2\kappa/a^2$, equivalent to Eq. (S8). We calculated the bending energy employing the MDAnalysis[9,10] MembraneCurvature library, with a bending modulus $\kappa$=10.

[7] H. A. Lorentz, Annalen der Physik **248**, 127–136 (1881).

[8] C. R. Company and W. H. Beyer, *CRC Standard Mathematical Tables: 28th Ed* (CRC Press, 1987).

[9] R. Gowers, M. Linke, J. Barnoud, T. Reddy, M. Melo, S. Seyler, J. Domański, D. Dotson, S. Buchoux, I. Kenney, and O. Beckstein, in *Proceedings of the 15th Python in Science Conference*, SciPy (SciPy, 2016) p. 98–105.

[10] N. Michaud-Agrawal, E. J. Denning, T. B. Woolf, and O. Beckstein, J. Comp. Chem. **32**, 2319–2327 (2011).
